# Toll Like Receptor 9 Pathway Mediates Schlafen^+^-MDSC Polarization During *Helicobacter*-Induced Gastric Metaplasias

**DOI:** 10.1101/2022.01.25.477562

**Authors:** Lin Ding, Jayati Chakrabarti, Sulaiman Sheriff, Qian Li, Hahn Nguyen Thi Hong, Ricky A Sontz, Zoe E Mendoza, Amanda Schreibeis, Michael A. Helmrath, Yana Zavros, Juanita L Merchant

## Abstract

**Background and Aims:** A subset of MDSCs that express murine Schlafen4 (SLFN4) or its human ortholog SLFN12L polarize in the *Helicobacter*-inflamed stomach coincident with intestinal or spasmolytic polypeptide-expressing metaplasia (SPEM). We propose that individuals with a more robust response to damage-activated molecular patterns (DAMPs) and increased Toll-like receptor (TLR9) expression are predisposed to the neoplastic complications of *Helicobacter* infection.

**Methods:** A mouse or human Transwell™ co-culture system comprised of dendritic cells (DCs), 2-dimensional gastric epithelial monolayers and *Helicobacter* were used to dissect the cellular source of interferon (IFNα) in the stomach by flow cytometry. Conditioned media from the cocultures polarized primary myeloid cells. Myeloid-derived suppressor cell (MDSC) activity was determined by T cell suppression assays. In human subjects with intestinal metaplasia or gastric cancer, the rs5743836 *TLR9*T>C variant was genotyped and linked to TLR9, IFNα and SLFN12L expression by immunohistochemistry. NFκB binding to the *TLR9* C allele was determined by electrophoretic mobility shift assays.

**Results:** *Helicobacter* infection induced gastric epithelial and plasmacytoid DC expression of TLR9 and IFNα. Co-culturing primary mouse or human cells with DCs and *Helicobacter* induced TLR9, IFNα secretion and SLFN^+^-MDSC polarization. Neutralizing IFNα *in vivo* mitigated *Helicobacter*-induced SPEM. The *TLR9* minor C allele creates an NFκb binding site associated with higher levels of TLR9, IFNα and SLFN12L in *Helicobacter*-infected stomachs that correlated with a greater incidence of metaplasias and cancer.

**Conclusion:** TLR9 plays an essential role in the production of IFNα and polarization of SLFN^+^-MDSCs upon *Helicobacter* infection. Subjects carrying the rs5743836 *TLR9* minor C allele are predisposed to neoplastic complications if chronically infected.

## Introduction

*Helicobacter pylori* (*H. pylori*) has infected more than half of the world’s population, of whom 5-15% develop gastric complications ranging from chronic atrophic gastritis and metaplasias to gastric adenocarcinoma (GAC). The first detectable host response to the pathogen is an increase in submucosal and intraepithelial inflammatory cells. Although inflammation is a protective response to fight the infection, immune suppressor cells create an environment favorable for hyperplasia, metaplasia and eventually tumor formation.

We previously reported that a subset of immune cells is recruited to the gastric epithelium during *Helicobacter* infection in mice and polarize into myeloid-derived suppressor cells (MDSCs)^1^. Polarization into MDSCs is highly correlated with the induction of Schlafen4 (SLFN4). SLFNs are a family of molecules, implicated in lymphoid and myeloid cell development and differentiation^2, 3^. In particular, SLFN4 is a myeloid cell differentiation factor that regulates myelopoiesis^4^. Similar to murine SLFN4, expression of its human ortholog SLFN12L increases in patients with gastric intestinal metaplasia (GIM) coincident with *H. pylori* infection and also marks a subpopulation of MDSCs^1^.

IFNα induces expression of all Schlafen family members^5–7^, which we recapitulated by treating primary murine myeloid cells with IFNα *in vitro*. Two months after *Helicobacter* infection, IFNα expression in the stomach increases with the induction of SLFN4 but prior to development of spasmolytic polypeptide expressing metaplasia (SPEM). Type I IFNs are important for the host’s defense against viruses, however, in bacterial infections, induction of these cytokines can lead to immunosuppression^8, 9^. Therefore, understanding the cellular source of IFNα during a *Helicobacter* infection might identify a potential therapeutic target that mediates the transition from chronic gastritis to GIM.

Although a variety of cell types produce type I IFNs, plasmacytoid DCs (pDCs) are considered the major source of type I IFNs in response to microbial ligands or modified host nucleic acids and proteins, mainly through endosomal sensors TLR7 or TLR9^10^. In particular, TLR9 recognizes hypomethylated CpG DNA, produced abundantly in *H. pylori*^11^.TLR9-mediated recognition of *H. pylori* DNA is a principal intracellular pathway induced by the live organism^12^. TLR9 activation recruits a complex comprised of MyD88, IL-1 receptor-associated kinases (IRAK4, IRAK1) and TNF-associated factor 6 (TRAF6) that ultimately induce expression of transcription factor interferon response factor 7 (IRF7)^13^. Expression of TLR9 increases in mouse gastric tissue following *H. pylori* infection^14^. Therefore, we posited that *Helicobacter* infection activates TLR9 in the stomach, triggering events that culminate in the local release of IFNα, SLFN4 expression and polarized SLFN4^+^ MDSCs. Indeed, enhanced TLR9 expression in the stomach as well as TLR9 polymorphisms have been linked to gastric cancer development^11, 15–17^. Mutations in the *TLR9* locus resulting in hyperactive TLR9 signaling have recently been reported for *Helicobacter*-infected subjects that developed GAC, underscoring the relevance of this pathway in the transformation process^15^.

Here we show that TLR9 plays an essential role in the production of IFNα and polarization of SLFN^+^-MDSCs in the stomach by co-culturing 2-dimensional mouse or human epithelial monolayers derived from gastric organoids with *Helicobacter spp*. and dendritic cells in the Transwell™ system. Neutralizing IFNα *in vivo* ameliorated

*Helicobacter*-induced gastric metaplasia. The C allele of the rs5743836 *TLR9* polymorphism creates an NFκB binding site and correlated with higher levels of TLR9, IFNα and SLFN12L protein in the stomachs *H. pylori*-infected subjects and a greater incidence of GIM or GAC in two independent Asian cohorts.

## Results

### *H. felis* infection induces gastric epithelial and pDC IFNα secretion

Both corpus and antral tissue levels of IFNα were significantly elevated after 2 mos of *H. felis* infection with only a slight increase in its serum level (Figure 1A). This result confirmed a local release of IFNα within the gastric mucosa as previously described^1^. IFNα was either expressed by cells PDCA-1^+^ pDC within the lamina propria, or by E-Cadherin^+^ gastric epithelial cells from infected mice (Figure 1B, C). Further characterization of the cellular source for IFNα was performed using flow cytometry (Figure 1D, E). Under uninfected conditions, EPCAM^+^ epithelial cells in both the gastric corpus and antrum constitutively expressed IFNα (Figure 1E). Following 2 mos of *H. felis* infection, there was a significant increase in IFNα-expressing cells, of which 45% were CD45^+^ cells infiltrating the corpus while about 26% infiltrated the antrum (Figure 1E). Most CD45^+^ cells (78%) appearing in the infected stomach were PDCA-1^+^CD11c^+^MHCII^+^CD11b^-^ plasmacytoid dendritic cells (pDCs) (Figure 1E). Although the total number of IFNα^+^ cells post infection was similar between the corpus and antrum (Figure 1F, G), pDCs were significantly higher in the corpus versus the antrum (Figure 1H). This possibly accounted for the different amounts of IFNα produced in these two compartments (Figure 1A), since pDC are known to secrete 10-100 times more type I IFNs (1-2 IU/cell) than other cell types in response to microbial stimuli^10^.

**Figure 1.**
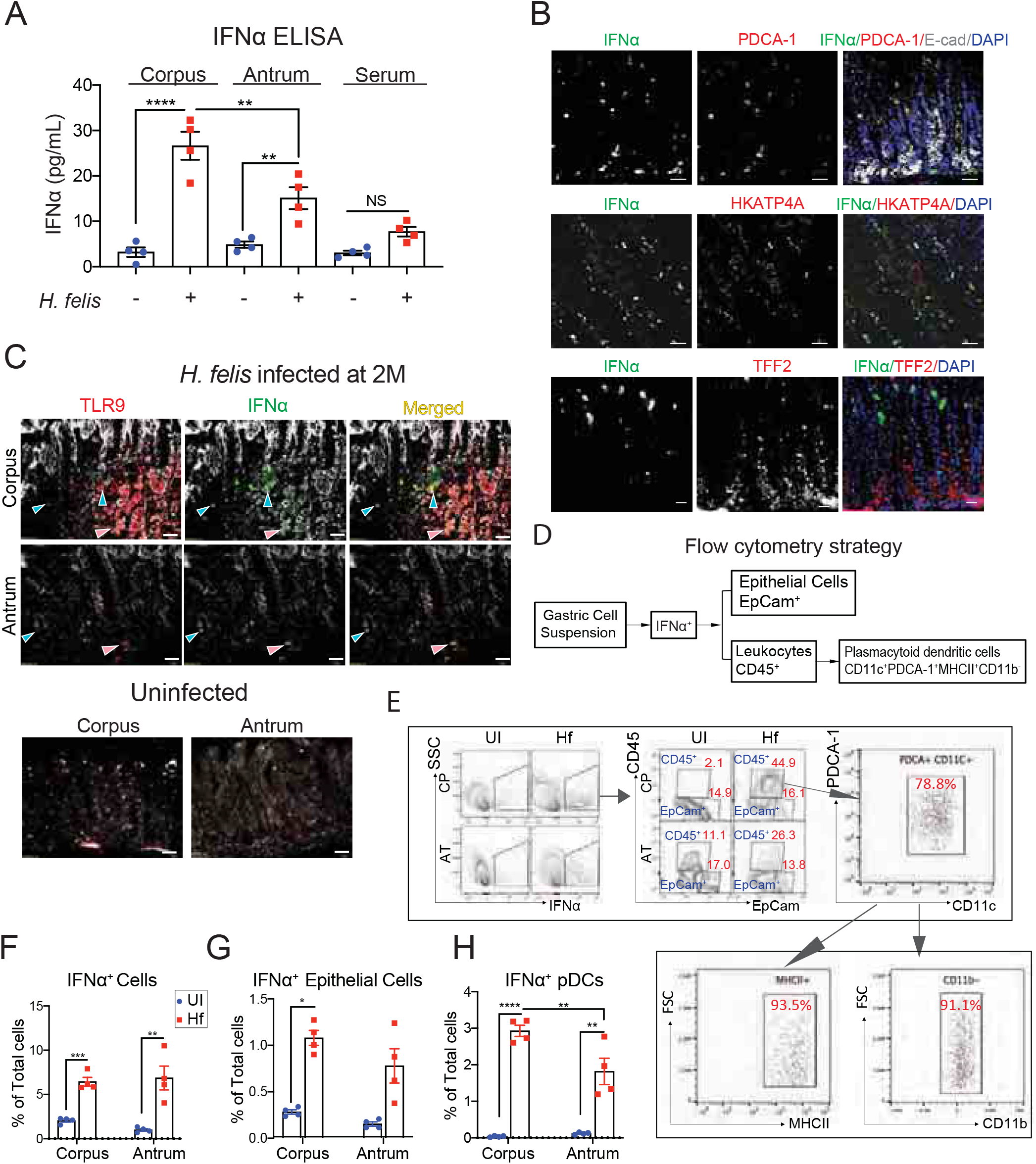
*H. felis* infection induced gastric epithelial and pDC expression of IFNα. Mice were infected with *H. felis* for 2 months. (A) IFNα levels in the corpus, antrum and serum from uninfected and infected mice were measured by ELISA. (B) Immunofluorescent images for IFNα (green), PDCA-1(red), E-cadherin (white) and TFF2 (red) in the gastric corpus of 2-month *H. felis*–infected mice. Scale bars: 10 μm. (C) Representative immunofluorescent images of gastric corpus and antrum immunostained for IFNα (green), TLR9 (red) and E-cadherin (white). Scale bars: 50 μm. Arrowheads indicate IFNα^+^TLR9^+^ cells within lamina propria (blue) and epithelial cells (pink), respectively. (D-E) Single cell suspensions from corpus or antrum were analyzed for IFNα, EpCam, CD45, PDCA-1, CD11c. Gated IFNα^+^CD45^+^PDCA-1^+^CD11c^+^ cells were further analyzed for MHCII and CD11b. Gated (F) IFNα^+^, (G) IFNα^+^ EpCam^+^ epithelial cells and (H) IFNα^+^ PDCA-1^+^CD11c^+^ pDC subpopulations as a percent of total number of gastric cell are shown in bar graphs. Shown are the median and interquartile range for N=4 mice per group. \**P*< .05. \*\**P*< .01. \*\*\**P*< .001. \*\*\*\**P*< .0001. N=4 mice per group.

TLR9 induces IFNα production^18^. Therefore, we further characterized the gastric cell populations expressing IFNα during a *Helicobacter* infection by immunofluorescence. The uninfected gastric epithelium weakly expressed TLR9 protein (Figure 1C). Two months after *H. felis* inoculation, IFNα colocalized with TLR9 in both epithelial (E-Cadherin^+^) and infiltrating immune cells. Thus, we concluded that *Helicobacter* infection induces TLR9 expression and IFNα production in both epithelial and immune cells (Figure 1C).

Since both epithelial cells and DCs produce IFNα, we compared their level of IFNα secretion with or without *H. felis* using a Transwell™ co-culture system comprised of pDCs and 2-dimensional (2-D) epithelial cell monolayers derived from either corpus organoids (CEM) or antral organoids (AEM) (Figure 2A). Confluent epithelial monolayers formed on collagen-coated Transwell™ inserts. Immunofluorescent staining demonstrated the presence of parietal (H^+^,K^+^-ATPase^+^), chief (pepsinogen C, PGC^+^), and surface mucous cells (Ulex Europaeus Agglutinin I, UEAI^+^) in the CEM, and endocrine cells (Chromogranin A^+^) including G cells (gastrin^+^) in the AEM (Figure 2B). PCR analysis using RNA collected from these monolayers confirmed the expression of parietal, surface mucous, chief and mucous neck markers in the CEM, as well as G cells, surface mucous cells and mucous neck cell markers in the AEM (Figure 2C), demonstrating that these epithelial monolayers expressed major cell lineages found in the native stomach.

**Figure 2.**
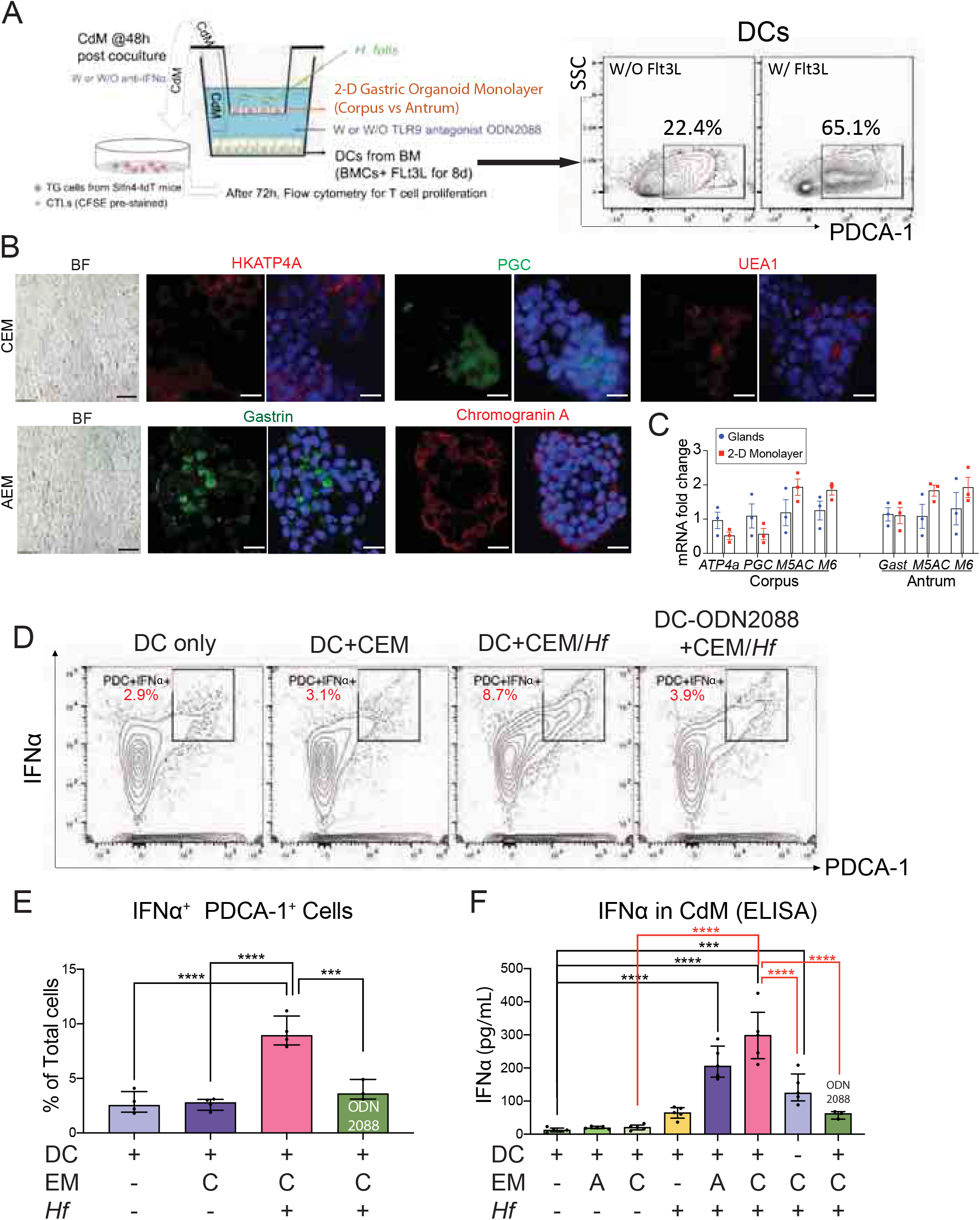
pDCs and epithelial cells secrete IFNα in the mouse co-cultures. (A) Schematic of the transwell system comprised of mouse pDCs, 2-dimensional (2-D) epithelial monolayers derived from corpus (CEM) versus antral (AEM) organoids cocultured with or without (MOI of 4) *H. felis*. pDCs were generated from mouse bone marrow cells by treating with recombinant Flt-3 ligand (~65% of these cells were PDCA-1^+^) seeded in the bottom of the Transwell. These polarized DCs were pre-incubated with or without TLR9 antagonist ODN2088. Live *H. felis* was then added to the upper Transwell chamber supporting the 2-D monolayers. Conditioned media (CdM) were collected from the bottom well 48 h after initiating the co-culture. A neutralizing antibody against IFNα was added to the CdM to identify the effect of IFNα. Thioglycollate-elicited peritoneal myeloid cells (TG cells) from Slfn4-tdTomato (Slfn4-tdT) mice were treated with CdM for 24 h, and then co-cultured with CFSE-pre-stained CTL cells for 72 h. (B) Brightfield (BF) and immunofluorescent (IF) staining of CEM vs AEM cultures demonstrating the expression of H^+^K^+^-ATPase (H, K-ATP4A, red), Pepsinogen C (PGC, green), surface mucous cells (UEAI, red) in CEM, and gastrin (green) and Chromogranin A (red) in AEM. Scale bar, BF 25μm, IF 50μm. (C) RT-PCR was performed using RNA extracted from mouse tissue and 2D monolayers quantify the expression of *Atp4a, PgC, Muc5ac (M5), Muc6* (*M6*) and *gastrin* (*Gast*) (bar graphs). (D) 48 h after co-culture, pDCs in the bottom well were analyzed for IFNα by flow cytometry. (E) Bar graphs show the quantitation. (F) An ELISA assay was performed to determine IFNα levels in the CdM. Horizontal lines represent the median and interquartile range. N=4 expts. \**P*< .05. \*\**P*< .01. \*\*\**P*< .001. \*\*\*\**P*< .0001.

pDCs were prepared from bone marrow-derived cells (BMDCs) treated with recombinant FLT-3 ligand to polarize ~65% of these BMDCs pDCs marked by PDCA-1^+^ (Figure 2A). pDCs were seeded in the bottom Transwell™ chamber and live *H. felis* was added to the upper chamber supporting the 2-D organoid monolayers (Figure 2A). The pDCs expressed higher levels of IFNα after co-culturing with *H. felis*-infected CEMs, but did not increase their expression in the absence of *H. felis* (Figure 2D, E). Moreover, IFNα induction was markedly reduced when the TLR9 antagonist ODN2088, which neutralizes the stimulatory effect of CpG ODNs, was added to the pDC culture (Figure 2D-F). To determine whether IFNα was present in conditioned media (CdM), an ELISA assay was performed using CdM collected from the wells 48 h after initiating the coculture with the bacteria (Figure 2F). Co-cultures with CEMs induced higher IFNα levels than AEMs, which was consistent with the in vivo results (compare Figure 2F to Figure 1A). Even without DCs, infected CEMs (CEM+*Hf*) released IFNα into the media, but to a lesser extent than when pDCs were present (DC+CEM+*Hf*) (Figure 2F). *H. felis* alone added to the upper chamber in the absence of epithelium produced some IFNα (DC+*Hf*), which was less than when the upper chamber contained *H. felis* co-cultured with the epithelial monolayers (DC+CEM+*Hf*) (Figure 2F). This result suggested that bacteria, their byproducts and soluble signals from the infected monolayers can diffuse through the 0.4μm pores to activate pDCs in the bottom chamber. Moreover, the TLR9 antagonist blocked the increase in IFNα suggesting that the secretion of this cytokine was due to TLR9 activation. Therefore, both epithelial and pDCs secreted IFNα, but secretion was most efficient when all 3 components were present.

Since Slfn4^+^-MDSCs are strongly induced by IFNα^1^, thioglycolate (TG)-elicited peritoneal myeloid cells were isolated from Slfn4-tdTomato (tdT) mice and treated with CdM for 24 h (Figure 3A, B). The epifluorescence from Slfn4-tdT^+^ cells increased after treating CdM from pDCs, gastric epithelial monolayers or both, only in the presence of *H. felis.* CdM from pDC/CEM/*H. felis* cocultures generated the highest number of Slfn4-tdT^+^ cells, which correlated with the CdM levels of IFNα (compare Figure 2F to Figure 3B). To test the function of IFNα in the CdM, a neutralizing antibody against IFNα was added to the CdM and blocked the appearance of Slfn4-tdT^+^ cells (Figure 3A, B). A CFSE-based T cell suppressor assay demonstrated that the presence of Slfn4-tdT^+^ cells activated by CdM from pDC/CEM/*H. felis* co-cultures suppressed T cell activation by ~50% (Figure 3C, D). Less T cell suppression (~25%) was observed using AEM-containing co-cultures. Moreover, the T cell suppressor function was blocked by IFNα antibody neutralization. Since TLR9 activation is a primary mechanism for IFNα induction, we showed that TLR9 expression was also induced by *H. felis* infection of both epithelial cells and pDCs (Supplementary Figure S1), along with induction of TLR9 signaling factors MyD88 and IRF7. Inhibiting TLR9 activation or neutralizing IFNα subsequently reduced SLFN4 induction and T cell suppression (Figure 3B,D). Therefore, we concluded that *H. felis* infection stimulates the release of IFNα from both epithelial cells and pDCs via TLR9 signaling, which induces *Slfn4* expression and polarization of myeloid cells to become SLFN4^+^-MDSCs, which suppress T cell proliferation.

**Figure 3.**
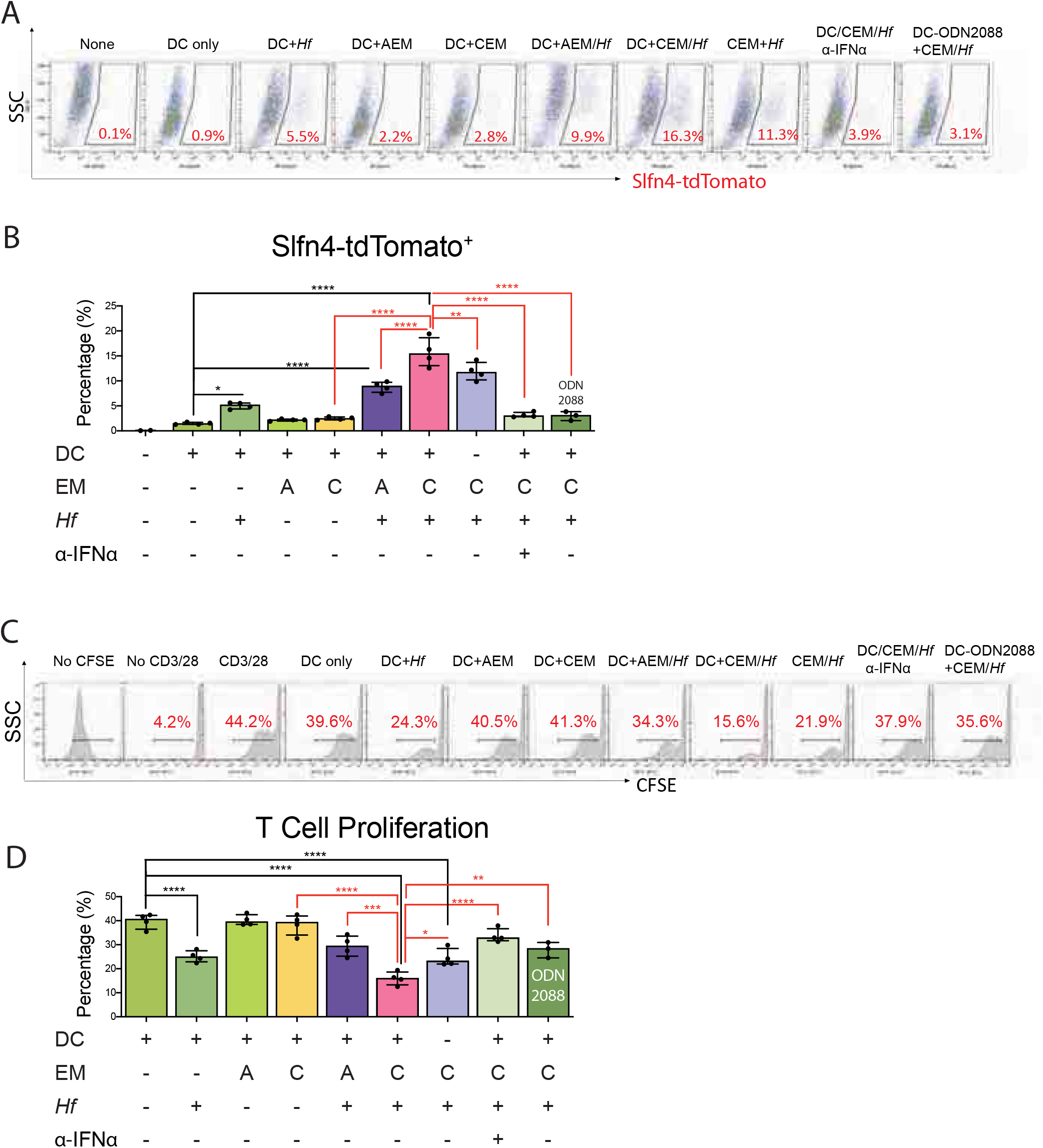
pDCs and epithelial cells induce Slfn4^+^-MDSCs via IFNα. Thioglycollate-elicited peritoneal myeloid cells (TG cells) from *Slfn4*-tdTomato (Slfn4-tdT) mice were treated with CdM from the 2-D monolayers/pDCs/*H. felis* co-culture (Figure 2) for 24 h, and then co-cultured with CFSE-prestained CTLs for 72 h. Flow cytometry was performed to analyze (A) the epifluorescence of Slfn4-tdT cells and (B) CFSE-based T cell suppression assay. The left three representative histograms in (B) show proliferation of control groups: No CFSE, without and with anti-CD3/28 microbeads to activate T cell proliferation. A neutralizing antibody against IFNα was added to the CdM to determine the effect of IFNα by flow cytometry (C) and quantified in the bar graphs (D). Horizontal lines represent the median and interquartile range for N=4 expts. \**P*< .05. \*\**P*< .01. \*\*\**P*< .001. \*\*\*\**P*< .0001.

### *H. pylori* polarizes human SLFN12L-MDSCs

To demonstrate that *H. pylori* induces human SLFN12L^+^-MDSCs, we performed a similar co-culture experiment using three separate human 2-D gastric organoid lines and immune cell populations generated from peripheral blood monocytes (PBMCs) (Figure 4A). The 2-D monolayers cultured in the upper Transwell™ chamber were treated with either wild type *H. pylori* (G27) or the *cag pathogenicity island* mutant (G27^ΔCag^)^19^. MDSCs, DCs and cytotoxic T cells (CTLs) were generated from PBMCs collected from the same patient. Consistent with SLFN4 induction in mice, SLFN12L was highly induced by CdM when the organoids were co-cultured with the G27 *H. pylori* wild type strain and less so by the G27^ΔCag^ mutant strain (Figure 4B, C). Moreover, SLFN12L induction by CdM was completely blocked by IFNα neutralizing antibody consistent with Schlafen regulation by type 1 IFNs. The IFNα-dependent increase in CD33^+^CD11b^+^-MDSCs was greatest when the cells were co-cultured with G27 but not with IFNα antibody (Figure 4D, E). TLR9 expression in the DCs was induced by G27 to a greater extent than by G27^ΔCag^ (Figure 4F, G), suggesting cooperation between epithelial cells and DCs, as observed in the mouse.

**Figure 4.**
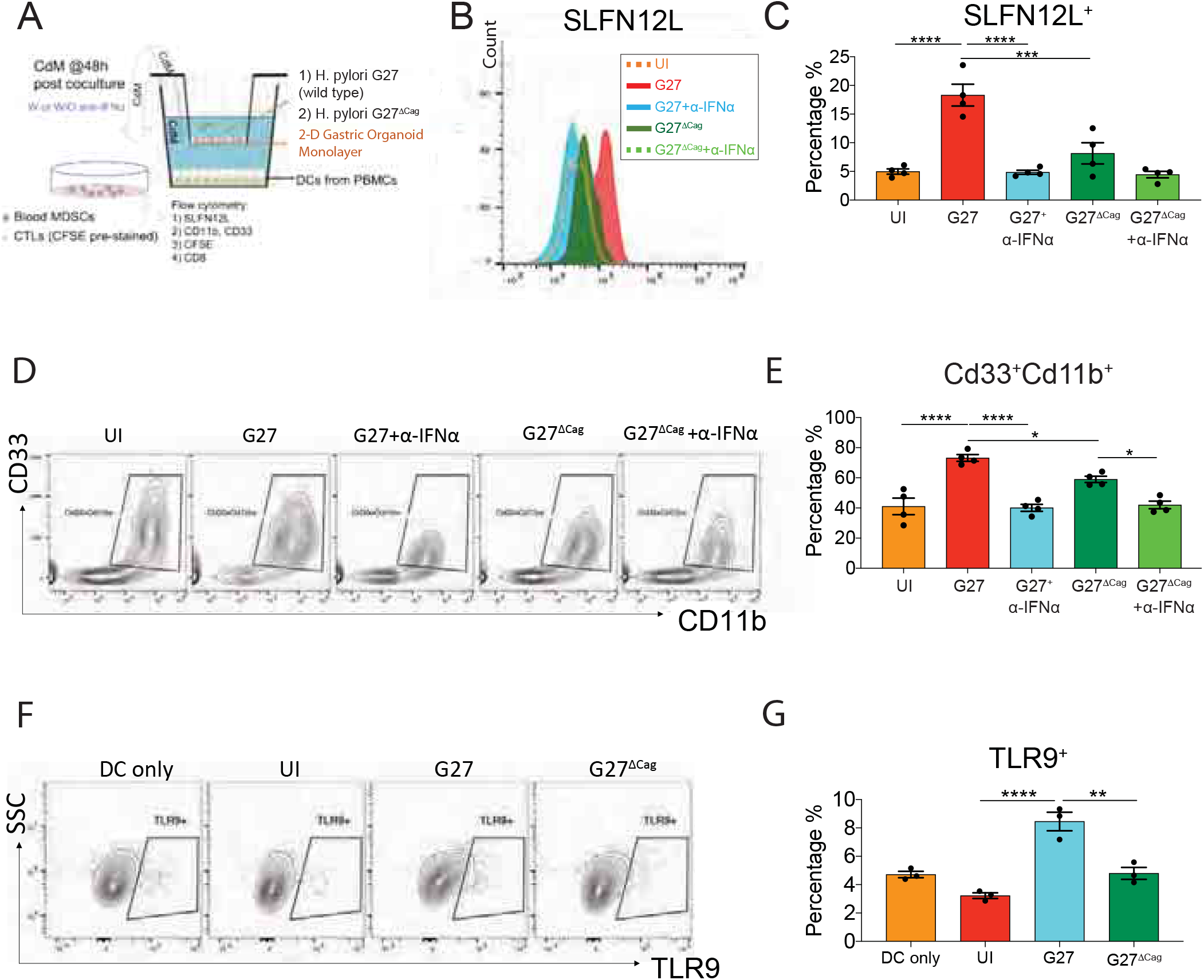
Human organoids/DCs/*H. pylori* co-culture experiment. (A) Schematic of the transwell system comprised of human DCs, 2-dimensional (2-D) corpus organoid-derived monolayers co-cultured with or without (UI) *H. pylori*. Dendritic cells were generated from PBMCs by treating with GMCSF seeded in the bottom of the transwell. Wild type *H. pylori* G27 strain and its CagA-mutant strain G27^ΔCag^ were added to the upper Transwell chamber supporting the 2-D organoid-derived monolayers. Conditioned media (CdM) were collected from the bottom well 48 h after initiating the co-culture. A neutralizing antibody against IFNα was added to the CdM to determine the effect of IFNα. SLFN12L^+^-MDSCs and CFSE-prestained CTLs from PBMCs were treated with CdM and then co-cultured together for 72 h. Flow cytometry was performed to analyze (B) SLFN12L and (D) Standard MDSC markers CD33, CD11b and (F) TLR9 in the DCs in the bottom of the well. (C, E, G) Bar graphs show the quantitation as a percentage of PBMCs. Horizontal lines represent the median and interquartile range for N=4 expts. \**P*< .05. \*\**P*< .01. \*\*\**P*< .001. \*\*\*\**P*< .0001.

### IFNα neutralization mitigates Helicobacter-induced SPEM

To further test the role of the TLR9/ IFNα/SLFN4 axis *in vivo*, we neutralized IFNα by injecting mice intraperitoneally (IP) with anti-IFNα antibodies (α-IFNα) (total of 72 μg per mouse × 3 injections) 4 mos after inoculating with *H. felis,* which reduced tissue IFNα levels (Figure 5A). The α-IFNα injections also reduced the elevated SLFN4 levels induced by the *Helicobacter* infection (Figure 5B). Four weeks after the α-IFNα treatments, less SPEM was observed as determined by significant recovery of parietal cells (H^+^,K^+^-ATPase4α) as well as a reduction in SPEM and associated markers TFF2 and Clusterin (Figure 5C). RT-PCR analysis confirmed less *clusterin, Tff2* and *CD44v9* mRNA and an increase in *H^+^,K^+^-Atpase4α* subunit *(Atp4a*) without significant recovery of the chief cell marker gastric intrinsic factor (*Gif*) (Figure 5D).

**Figure 5.**
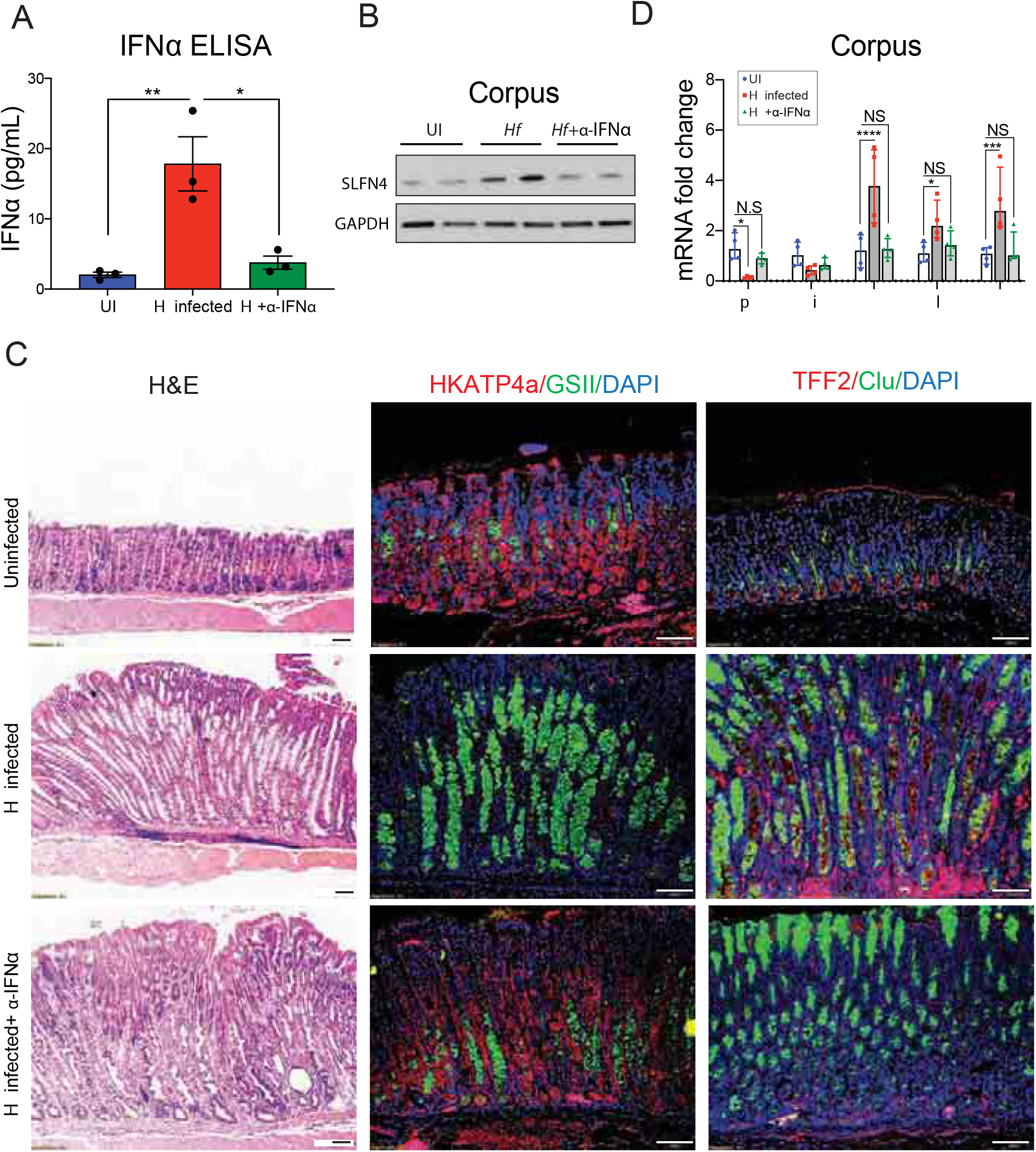
Neutralizing IFNα in vivo during *H. felis* infection reduces SPEM. (A) IFNα levels in the corpus from uninfected, infected and IFNα neutralized mice were measured by ELISA. (B) Western blot of SLFN4 expression in mouse corpus, GAPDH as a loading control. (C) Histologic changes were shown by H&E, H^+^,K^+^-ATP4α (green), GSII (green), TFF2 (red) and Clusterin (green) IF. Scale bar, 50μm. (D) SPEM markers *ATP4α, Gastric intrinsic factor (Gif), Tff2, Clusterin (Clu)* and *CD44v9* mRNA were analyzed by qPCR. Horizontal lines represent the median and interquartile range for N=3 mice per group. One-way ANOVA followed by Tukey’s multiple comparisons test on log-transformed values is shown. \**P*< .05. \*\*\**P*< .001. \*\*\*\**P*< .0001. NS, not significant.

### rs5743836 *TLR9 C allele* correlates with *H. pylori*-induced gastric lesions

Since TLR9 plays an essential role in the induction of IFNα by *H. pylori*, and enhanced *TLR9* expression in the stomach as well as *TLR9* polymorphisms have been linked to gastric cancer development^11, 15^, we examined the association of two *TLR9* single nucleotide polymorphisms (SNPs) (rs5743836*TLR9*T>C at-1237 and rs187084*TLR9*C>T at-1486) residing 249 nucleotides apart within the *TLR9* promoter^20^. Therefore the presence of these two SNPs was examined in two previously characterized Asian cohorts with gastritis and IM or GAC^21^. Subjects with atrophy and IM were associated with a higher frequency of the rs5743836 minor C allele than patients with just gastritis (Figure 6A). Moreover, analysis of a Chinese cohort with GAC similarly showed a greater association of the TLR9 C allele with GAC compared to subjects without cancer (Figure 6A). The TLR9 rs187084 SNP at-1486 showed no correlation with the histologic lesions (Supplementary Table S1).

**Figure 6.**
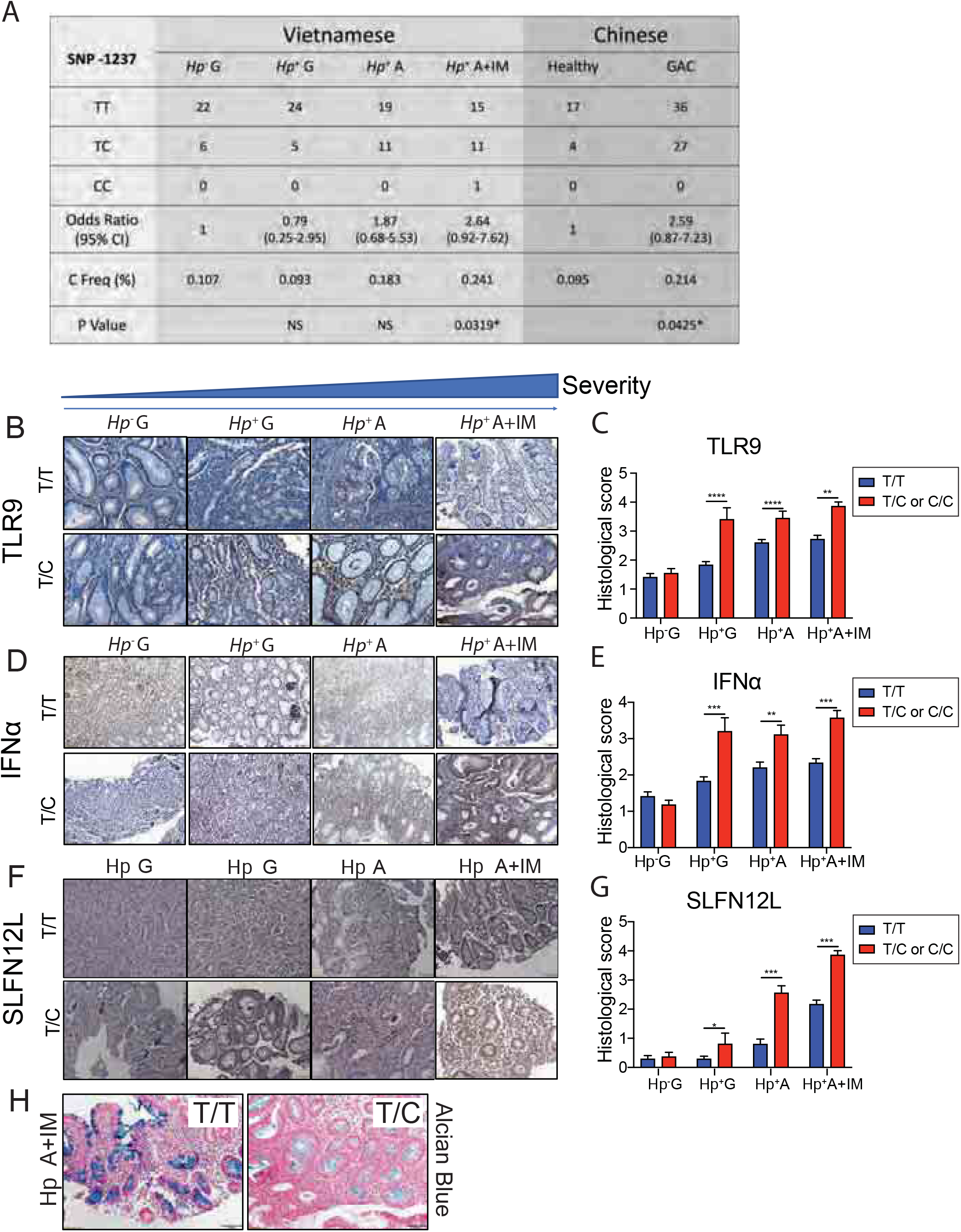
The rs5743836 TLR9T>C SNP correlated with IM, GAC and higher TLR9/IFNα/SLFN12L expression in the infected stomach. (A) Association of the minor TLR9 C allele (at-1237 from promoter start site) frequency with gastric metaplasia or gastric cancer (GAC) in Vietnamese and Chinese patient cohorts. * *P*< .05 over uninfected control group within the cohort. NS, not significant. *HP: H. pylori*; G: Gastritis; A:gastric atrophy; IM:Intestinal metaplasia; GAC:gastric cancer. Representative images showing IHC staining for (B) TLR9 (D) IFNα and (F) SLFN12L in gastric biopsies from uninfected Vietnamese patients with gastritis (*Hp*^-^ G), *H. pylori*-infected gastritis (*Hp*^+^ G), *H. pylori*-infected patients with gastric atrophy (*Hp*^+^ A) or *H. pylori-infected* patients with atrophy and IM (*Hp*^+^ A+IM) carrying both major T alleles *TLR9*-1237T/T or carrying at least one minor *TLR9*-1237C allele. Mag: 200X. Histologic scoring of (C) TLR9 (E) IFNα and (G) SLFN12L were shown as bar graphs. Kruskal-Wallis ANOVA with Dunn’s test of multiple comparisons was performed. (H) Representative Alcian blue staining for patients with intestinal metaplasia and atrophy and carrying either the T/T or T/C alleles. Horizontal lines represent the median and interquartile range. \**P*< .05. \*\**P*< .01. \*\*\**P*< .001. \*\*\*\**P*< .0001.

We used immunohistochemistry to determine whether the rs5743836 minor allele correlated with increased expression of TLR9, IFNα, and SLFN12L in gastric biopsies from the Vietnamese cohort. In the patients with gastritis but negative for active *H. pylori* (CLO-), there was minimal expression of TLR9 in the gastric epithelium and lamina propria. By contrast, subjects with atrophy or IM showed elevated TLR9 expression if heterozygous for the rs5743836 C allele compared to those subjects homozygous for the major T allele (Figure 6B, C). IFNα expression primarily within the lamina propria was also higher in subjects carrying at least one C allele (Figure 6D, E). Moreover, SLFN12L expression was higher in the lamina propria of *H. pylori*-infected gastric mucosa with atrophy and IM (Figure 6F, G). Representative Alcian blue staining for patients with the T/T allele and the T/C allele is shown (Figure 6H). Therefore, *H. pylori*-infected subjects carrying the rs5743836 C variant tended to express higher levels of TLR9, IFNα and SLFN12L.

### rs5743836 TLR9 C allele correlated with higher NFκB binding activity

In silico analysis revealed that rs5743836 TLR9 creates a putative NFκB binding site at-1237, which we confirmed by EMSA. TNFα induced expression of nuclear NFκB within 30 min (Figure 7A). NFκB binding to the probe containing the *TLR9*-1237C variant was detected in the absence of cytokine (Con) but was highly induced by TNFα (Figure 7B). Binding specificity was confirmed by competitive displacement with 200-fold excess of unlabeled probe. In contrast, no specific nuclear protein binding from either control or the TNFα-treated group was observed using the *TLR9*-1237T allele (Probe T) (Figure 7B). The EMSA complex binding to the minor C allele was completely disrupting and supershifted with NFκB antibody (Figure 7C). Therefore, the rs5743836 TLR9 minor C allele forms an NFκB binding site, which leads to higher levels of NFκB binding, and expectantly higher *TLR9* expression.

**Figure 7.**
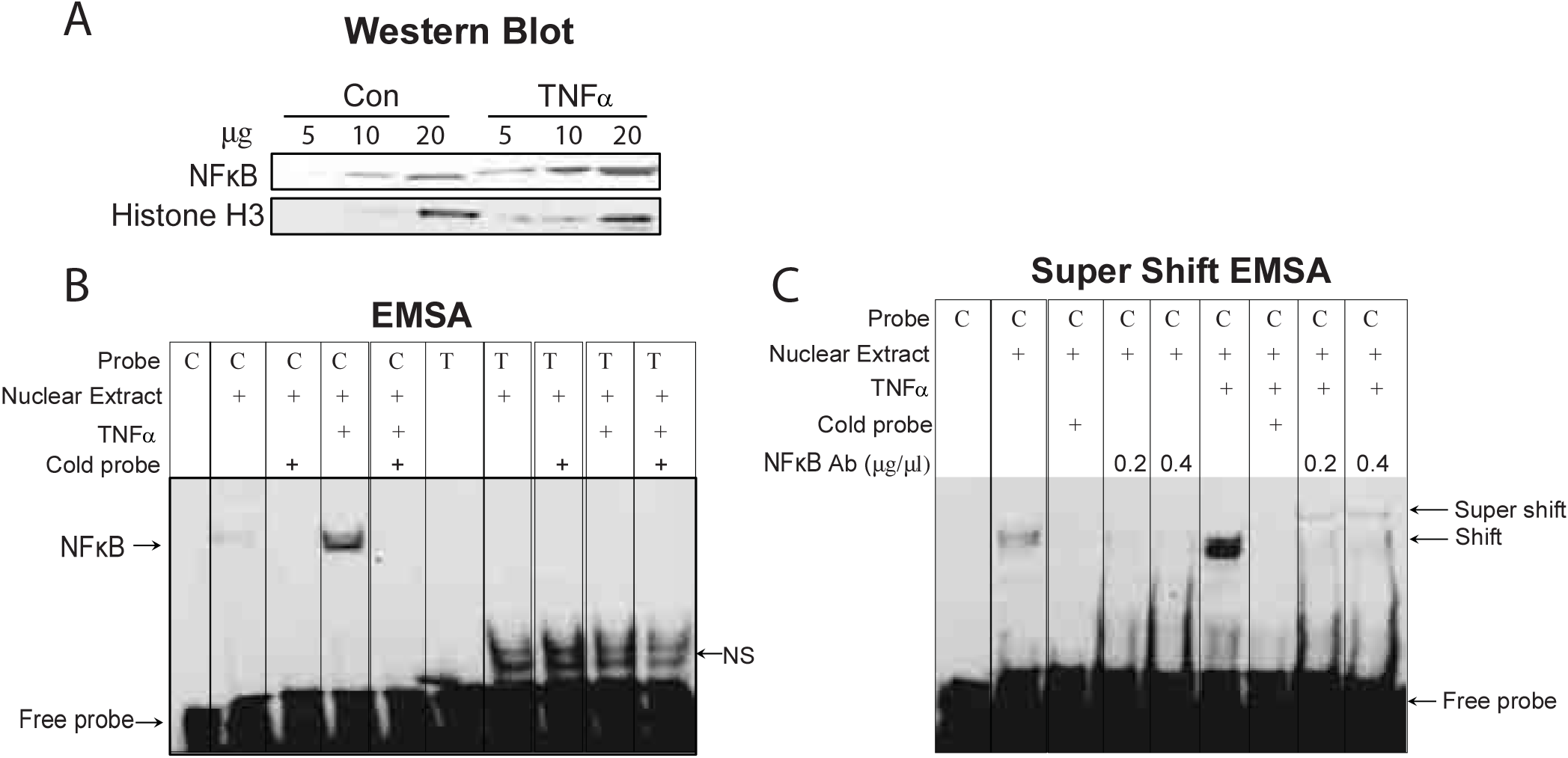
NFKB binding activity correlated with rs5743836TLR9C allele. HEK-293 cells were treated with PBS (Con) or TNFα (30ng/mL) for 30 min to induce NFκB prior to extracting nuclear proteins. (A) A western blot was performed to confirm NFκB expression in 5, 10, 20 μg of nuclear extract. Histone H3 was the loading control. (B) Electrophoretic mobility shift assays (EMSA) were performed to demonstrate NFκB DNA binding to the double-stranded probe consisting of only the major T allele *TLR9* - 1237T (Probe T) versus the minor C allele −1237C (Probe C). NFκB, nonspecific binding (NS), and free probe are indicated by arrows. 200-fold excess of unlabeled (cold) probe was used as competitor. (C) Two concentrations of NFκB antibody (0.2; 0.4 μg/μL) was used to disrupt and supershift the protein bound to Probe C.

## Discussion

*Helicobacter pylori* infection predisposes to lifelong chronic gastric inflammation, significantly increasing an individual’s risk for GAC. Immune suppressor cells dampen the active inflammatory process, which hastens tissue repair upon resolution of the infection^22^. However, a byproduct of the mucosal repair process is release of proliferative signals that create a permissive microenvironment for hyperplasia, metaplasias, and eventually neoplastic transformation ^23^. A subset of MDSC immune suppressor cells expresses members of the Schlafen gene family--murine Schlafen4 (SLFN4) or its human ortholog SLFN12L. We previously reported that Schlafens are induced in the *Helicobacter*-infected stomach coincident with atrophy and the appearance of gastric metaplasias, i.e., SPEM in mice and IM in humans^1^. With gastric atrophy, the mix of cellular and bacterial debris generated by the chronic infection releases DAMP ligands from stressed or dying cells that activate intracellular TLR9. Thus, release of DAMPs into the extracellular milieu is one mechanism by which inflammation creates an environment favoring cancer development^24^. TLR9 activation initiates a cascade of IFN-regulated molecules, e.g., IRFs, Schlafens^25^. Here we report that both *Helicobacter*-infected pDCs and gastric epithelial cells produce IFNα in response to TLR9 activation. IFNα is required for Schlafen expression and subsequent polarization of SLFN^+^-MDSCs that occur during the transition from an acute to a chronic *Helicobacter*-induced infection ^1, 25^.

Emerging studies have shown that type 1 IFNs potently regulate both innate and adaptive immune responses ^26^ and mediate pro-inflammatory and immunomodulatory functions^27^. During an infection, DCs migrate to the epithelial layer and intercalate into the epithelium to sample the *H. pylori*-colonized lumen^28^. Here we report that after 2 months of a *Helicobacter* infection, IFNα was expressed by both epithelial cells and infiltrating immune cells within the lamina propria, many of which were pDCs ^12, 29^. Once activated, pDCs secrete copious amounts of type I IFNs via TLR9-MyD88-IRF7 ^29, 30^. inflammasome signaling TLR9 activation by *H. pylori* immunomodulatory DNA sequences can enhance *H. pylori* chronicity by suppressing pro-inflammatory cells^11,14,28^. In addition, the CpG-DNA-binding protein high-mobility group box 1 (HMGB1) and mitochondrial DNA are epithelial-derived sources of DAMP ligands known to bind to TLR9 ^31,32^. Since acid secreting parietal cells are rich sources of mitochondria, this might explain why the corpus epithelium exhibited stronger IFNα induction. Indeed, we previously showed that co-expression of the apoptotic marker caspase-3 and IFNβ occurs on parietal cells at 2 months post infection, demonstrating that dying parietal cells are at least one potential source of DAMP ligands^1^.

Examination of a frequently studied TLR9 SNP in the promoter that creates an NFκb binding site demonstrated an increased risk of GIM and GAC. This result underscores the strong influence of inducing and maintaining elevated levels of NFκB in mediating gastric tumor cell growth^21^. Although it is not understood why some individuals are predisposed to the neoplastic consequences of chronic *H. pylori* infection, linkage to this particular *TLR9* SNP that favors greater NFκB binding implicates a genetic bias for some patients to develop neoplasia. Identifying the rs5743836 TLR9 minor C allele in different ethnic populations might provide greater understanding of who is susceptible to the more serious complications from chronic *H. pylori* infection and therefore require increased endoscopic surveillance.

With respect to treatment options, it is controversial whether TLR9 CpG ODN agonists induce the accumulation of tumor-infiltrating MDSC, e.g., in pancreatic ductal adenocarcinoma^33^; while other studies suggest that CpG DNA is essential for decreasing MDSC suppressive activity^34, 35^. Therefore, the regulation of MDSC polarization by increased TLR9 activity might have contradictory effects depending on the source and quantity of the DAMP ligands. Several TLR9 DNA agonists used as adjuvants for vaccines against cancer or infectious diseases are in clinical trials, although none are being tested on gastric cancer ^36^. In summary, the current study suggests that TLR9 activation and subsequent IFNα production helps to shape the immune microenvironment preceding tumor development and that suppression of DAMP pathway components might ultimately be therapeutic targets to consider for cancer monotherapy or co-therapy^37^.

## Materials and Methods

### Patient samples

The two TLR9 SNPs were genotyped by direct DNA sequencing. TLR9, IFNα and SLFN12L expression were demonstrated by immunohistochemistry (IHC) in two previously described cohorts of de-identified Asian subjects^21^. Eighty-four subjects were from Xiangya Hospital (China) (IRBMED; ID: HUM00113773), containing 21 healthy controls and 63 patients with gastric cancer. One hundred and fourteen subjects were from the Institute of Biotechnology in Vietnam (IRBMED; ID: HUM00108090), containing 28 gastritis cases without an active *H. pylori* infection, 29 cases with an active infection, 30 with gastric atrophy and 27 with gastric atrophy/IM. Detection of *H. pylori* infection was performed using *H. pylori* IgG ELISA and the Campylobacter-Like Organism (CLO) test at endoscopy. A GI pathologist blinded to the clinical diagnosis reviewed the biopsies. Human-derived gastric organoids, MDSC, CTLs and DCs were generated from corpus tissue and autologous whole blood was obtained from patients undergoing sleeve gastrectomy at Cincinnati Childrens’ Hospital and Medical Center (IRB ID: 1912208231).

### Histological analysis

Five-micron paraffin sections were prepared, de-paraffinized in a xylene-alcohol series, followed by rehydration in phosphate-buffered saline (PBS). Antigen retrieval was performed using 10 mM citric acid (pH 6). Slides were washed twice in 0.01% Triton X-100 in PBS, incubated for 1 h with the serum of the animal in which the secondary antibody was raised. IHC was performed using the Vectastain ABC kit (Vector Laboratories) and DAB substrate kit (ab64238, Abcam). Primary antibodies were TLR9 (1:100, Abcam, #ab134368), IFNα (1:100, Abcam, #ab7373) and SLFN12L (1:100, NBP1-91060, Novus). Secondary antibodies were goat anti-rabbit (1:1000, #656140, Invitrogen) and donkey anti-mouse (1:500, BA9200, Vector Laboratories). Immunofluorescence (IF) was visualized using an Olympus BX53F microscope (Center Valley, PA). Two researchers blinded to the experimental conditions used a semiquantitative scale from 0 to 4 on 200x histologic fields to score TLR9/IFNα/SLFN12L expression (0=negative; 1 = weak positive; 2 = intermediate; 3 = strong; 4= very strong). The weighted H score = (% cells with a score = 1) + 2× (% cells with a score = 2) + 3× (% cells with a score = 3) + 4× (% cells with a score = 4). For each slide, 6-10 microscopic fields were scored.

Specific protein expression was identified by IF. Antigen retrieval was performed paraffin sections using 10 mM citric acid (pH 6). For 2-D epithelial monolayers, Transwell™ insert membranes were cut out and fixed in 3.7% formaldehyde for 10 min. Slides or membranes were washed twice in 0.01% Triton X-100 in PBS, incubated for 1 h with the serum of the animal in which the secondary antibody was raised and then incubated with the following antibodies overnight: Lectin GSII (Fisher, # L21415), ATP4a (Invitrogen, #MA3-923), Clusterin (Abcam, # ab184100), trefoil factor 2 (TFF2, Abcam, #ab49536), Pepsinogen C (Abcam, #ab255826), Ulex Europaeus Agglutinin I (UEA, Novus, # NB110-13922), Gastrin (DAKO, #A0568), Chromogranin A (Abcam, #ab17064). Primary antibodies were detected using Alexa Fluor-conjugated secondary antibodies (Molecular Probes, Invitrogen) at a dilution of 1:500. Slides were mounted with Prolong gold anti-fade reagent containing DAPI (Life technologies). IF was visualized using the Olympus BX53F microscope.

### 2-D Gastric Epithelial Monolayer/Helicobacter/Dendritic cell co-culture (mouse)

Before the co-culture experiment, plasmacytoid dendritic cells (pDCs), 2-D gastric epithelial monolayers derived from organoids and *H. felis* were prepared separately (Figure 2A).

Plasmacytoid DCs (pDCs): Bone marrow (BM) cells (10^6^) collected from the hind leg long bones of 8 to 12-week-old mice were suspended in the RPMI 1640 media with 10% FCS and then seeded in 24-well plates. pDCs were generated by treating BM cells with 200ng/ml Flt-3 ligand (Abcam, ab270071) at 37°C for 8 days ^39^. DCs were preincubated with or without the TLR9 antagonist ODN2088 (5μM, InvivoGen, # tlrl-2088) for 1.5 h before co-culture and throughout the co-culture.

Gastric epithelial monolayer: Mouse gastric organoids were generated from the corpus or antrum and cultured in Matrigel at 37°C in 5% CO_2_ as previously described^21^. After 3-D gastric organoids were generated for 6-7 days, they began to transition into 2-D epithelial monolayers. Transwell inserts (6.5 mm diameter, 0.4 μm pore size, Corning) were uniformly coated with Collagen I (Sigma Aldrich, C3867) to form thin (<20 μm) films, by spreading the substrate over the insert surface using 500 μL of ice cold 50 μg/mL Collagen I followed by an incubation at 4 °C overnight. Organoids were harvested in cold advanced DMEM/F12 by pipetting up and down and then were transferred into 5mL round-bottom tubes followed by the centrifugation at 200x g for 5 min at 4°C. Pellets were re-suspended in 1mL/tube of pre-warmed (37°C) 2-D gastric monolayer growth media (Advanced DMEM/F12 medium supplemented with 2mM L-glutamine, 10 mM Penicillin/Streptomycin, 10% Fetal Calf Serum, 10 mM HEPES Buffer, 1 x N2, 1 x B27, supplemented with 10 nM gastrin, 50 ng/mL Epidermal Growth Factor, 1 μM TGF-β inhibitor, and 10 μM Y-27632 ROCK inhibitor) and then seeded onto the collagen coated Transwell™ substrates. Plates with Transwell™ inserts were incubated at 37 °C for 4 days to form the monolayer with >90% confluence (Figure 2B) ^38^.

The *H. felis* CS1 strain was cultured in sterile-filtered Brucella broth (BD) plus 10% horse serum (Atlanta Biologicals) using the GasPak EZ Campy Container System (BD Biosciences) at 37°C with shaking at 100 rpm.

*H. felis* was resuspended in 2-D monolayer growth media and then added to the inserts containing the monolayers at a multiplicity of infection (MOI) of 4. The inserts were then combined with the plates containing pDCs in the bottom well to initiate the coculture^38^. After 48h of co-culture, the conditioned media (CdM) was collected and treated with or without IFNα (Abcam, #ab7373) neutralization antibody to polarize MDSCs and test their activity in a T cell proliferation assay.

### 2-D Gastric Epithelial Monolayer/Helicobacter/Dendritic cell co-culture (human)

DCs, 2-D gastric organoid-derived monolayer and *H. pylori* were cultured separately ^38^ (Figure 4A). Gastric organoid were generated from gastric biopsies collected from 4 patients undergoing sleeve gastrectomy ^38^. The 2-D organoid-derived monolayers plated in the upper chamber of the Transwell were infected with 0.1-0.2 million wild type *H. pylori* (G27) with an intact *cag* PAI encoding the type IV secretory system (T4SS) or the *cag* PAI mutant strain (G27^ΔCag^), which is unable to synthesize CagA protein preventing injection of bacterial DNA and virulence factors into host cells.

MDSCs, DCs and cytotoxic T cells (CTLs) immune populations were generated from peripheral blood monocytes (PBMCs) collected from the same patient. Generating the monolayer and DC co-culture was as described in the mouse co-culture experiments. After 48h co-culture, the cell media (CdM) was collected and treated with or without IFNα neutralization antibody (PBL Assay Sciences, #2112-1) prior to addition to MDSCs. Human MDSCs and CTLs were isolated from PBMCs ^38^. CTLs were activated using anti-CD3/CD28^+^ beads (ThermoFisher, #11161D), cultured overnight, pre-labeled with 5 μM CFSE at 37 °C for 20 min, and then co-cultured with MDSCs for 72 h.

### Flow cytometry

A single cell suspension was isolated from the mouse gastric corpus or antrum ^1^. Mouse DCs were harvested by scraping the well with a tissue culture scraper and then rinsing with PBS. The cells were stained with EPCam (Biolegend # 118230), CD45 (Biolegend # 103146), CD11c (ThermoFisher # 46-3697-82), PDCA-1 (Biolegend, #127103), IFNα (Abcam, #ab7373), CD11b (Biolegend #101239), MHCII (ThermoFisher # 56-5321-82). Human DCs, MDSCs and CTLs in the dish were harvested in PBS and stained with SLFN12L (Novus Biological NBP 1-91060), CD33 (Biolegend # 303428), CD11b (Biolegend # 301308), PDCA-1 (ThermoFisher # 53-3179-4), TLR9 (Biolegend #394803). Zombie dye (Biolegend # 423107) was used for live cell gating. Compensation beads (#01-2222-41) and ArC™ Amine Reactive Compensation Bead Kit (#10628) were purchased from Thermo Fisher. The cells were analyzed using a Cytek Aurora 5 Laser Spectral Flow Cytometer (Sony Biotechnology).

### Treatment with a monoclonal anti–IFNα antibodies

*pCMV-Shh transgenic* and wild type mice on a C57BL/6 background^1^ were infected with *H. felis* for 4-4.5 mos and then were injected intraperitoneally three times with 1.2 mg/kg of a monoclonal anti-IFNα antibody (Abcam, #ab7373) spaced three days apart for each administration for a total of 72 μg. Mice not receiving anti-IFNα antibody received saline. The mice were necropsied four weeks after the first injection.

### Statistical Analysis

For ELISA, statistical analysis for significance was performed on the log-transformed values using one-way analysis of variance with Tukey’s posthoc test for multiple comparisons (GraphPad Prism). Cell subpopulations were computed as percentages and the weighted scores used in the morphometric analysis were analyzed using Kruskal-Wallis one-way ANOVA with Dunn’s test for multiple comparisons (GraphPad Prism). The quantitation of western blots by ImageJ reflected the amount as a ratio of each protein band relative to the lane’s loading control from 3 experiments. Hardy-Weinberg equilibrium of alleles was assessed using Chi-squared analysis. The association between the polymorphism and gastric lesion risk was estimated using an odds ratio (OR) and a 95% confidence interval (CI). All data were expressed as the median with the Interquartile range (IQR). P<0.05 were considered statistically significant. The number of samples per group and replicate experiments are indicated in the figure legends.

## Declaration of Interests

The authors have declared that no competing interests exist.

## Author Contributions

LD designed research studies, conducted experiments, analyzed data, and wrote the manuscript. JC and YZ performed human cell co-culture experiments. SS performed EMSA and western blot analysis. QL, HH, AS and MH collected and processed clinical samples. RS performed animal breeding, husbandry and treatments. ZM performed IHC staining and analysis. JLM designed research studies reviewed all data and wrote the manuscript.

## Acknowledgements

The authors thank Olivia Q. Merchant for assistance with figure layouts.

## Abbreviations

DAMP: Damage-associated molecular patterns
pDC: plasmacytoid dendritic cells
GAC: gastric adenocarcinoma
SPEM: spasmolytic polypeptide-expressing metaplasia
IM: intestinal metaplasia
MDSC: myeloid derived suppress cells

## Supplementary Methods

### Genotyping of TLR9 single nucleotide polymorphisms

Genomic DNA samples were extracted from paraffin blocks using the QIAamp DNA FFPE Tissue Kit (Qiagen). Two TLR9 single nucleotide polymorphisms (SNPs) (rs5743836T>C at −1237and rs187084C>T at −1486) were identified by direct nucleotide sequencing of PCR products. Forward primer −1486/1237F: 5’-TCCCAGCAGCAACAATTCATTA-3’ and reverse primer −1486/1237R: 5’-CTGCTTGCAGTTGACTGTGT-3’. The PCR conditions used were as follows: One cycle at 95°C for 5 min, 36 cycles of denaturation at 94°C for 40 s, annealing at 58°C for 40s, extension at 72°C for 1 min, and a final extension step at 72°C for 10 min. The sequencing was carried out using the 3730 DNA Analyzer Applied Biosystems with the sequencing primer −1486/1237S: 5’-GGGTGTACATAATTCAGCAG-3’.

### T cell suppression assay

Thioglycollate-elicited peritoneal myeloid cells (TG cells) isolated from Slfn4-tdTomato mice as described previously^1^ were treated with the CdM from 2-D Gastric epithelial monolayer/*Helicobacter*/Dendritic cell co-culture at 37°C for 24 h. Induction of Slfn4-tdT^+^ cells was analyzed by the FACS Canto II cell analyzer (BD). T cells from the mouse spleen were isolated using the EasyStep Mouse T Cell Isolation Kit (StemCell Technologies) and pre-stained with carboxyfluorescein diacetate succinimidyl ester (CFSE) using the CellTrace CFSE Cell Proliferation Kit (Molecular Probes, #C34554). T cells were cultured in RPMI 1640 supplemented with 10% FCS, 2 mM L-glutamine, 1 mM sodium pyruvate, 100 mM nonessential amino acids, 5 mM HEPES free acid, 10 ml of 0.05 μM β-mercaptoethanol, and 100 U/mL penicillin/streptomycin with 10% heat-inactivated FBS. To stimulate proliferation, the T cells were cultured with anti-CD3/CD28–coated sulfate latex beads added to the media. Suppression of T cell proliferation was assayed after the addition of treated TG myeloid cells as described above for 3 days at a T cell/SLFN4^+^ cell ratio of 10:1. Cell proliferation was analyzed using the FACS Canto II cell analyzer (BD).

### IFNα ELISA

Mouse serum was obtained by centrifugation at 2000 rpm for 30 min at room temperature. Protein was extracted by homogenizing gastric tissue in RIPA buffer supplemented with proteinase inhibitor (ThermoFisher, #78425) and the supernatant was used in the assay. IFNα levels were measured using the Mouse IFN Alpha All Subtype ELISA Kit (PBL, #42115-1), per manufacturer’s instructions.

### Western Blot Analysis

Protein extracts from DCs in the bottom Transwell were collected in RIPA buffer after co-culture (Pierce, Rockford, IL). Tissue from the corpus was homogenized in RIPA buffer. Two-dimensional gastric epithelial monolayers were scraped off the Transwell inserts and the protein was extracted in RIPA buffer. Nuclear extracts from HEK-293 cells were cultured and extracted as described below for EMSA. Western blots were performed by re-suspending proteins in sample buffer, and then resolving on Novex Tris-Glycine 4-20% gradient gels (Invitrogen) with Tris running buffer; transferred to the PVDF membrane using the iBlot Dry Blotting System (Invitrogen) according to the manufacturer’s instructions. The membranes were blocked in 5% nonfat milk for 1 h at room temperature and then sequentially probed overnight at 4°C with primary antibodies to TLR9 (Abcam, #134368), MyD88 (Santa Cruz, #sc74532), IRF7 (Santa Cruz, #74471), β-Tubulin (Cell signaling, #5346S), NFκB (Cell signaling, #8242), SLFN4 (custom made by GenScript), GAPDH (MA5-15738, Thermo Fisher) or Histone H3 (Cell signaling, #12648).

### Electrophoretic Mobility Shift Assay (EMSA)

HEK-293 cells were grown in DMEM-10% FBS media. Cells were serum starved in 0.5% FBS for 16h until 70% confluent. The next day, cells were treated with 0.1% BSA in PBS or human TNFα (30 ng/mL for 30 min) in 0.1% BSA in PBS and then washed in PBS. Nuclear extracts were prepared using the NE-PER Nuclear and Cytoplasmic Extraction Kit (# 78833, ThermoFisher). EMSAs were performed using the LightShiftR Chemiluminescent EMSA Kit (#20148, ThermoFishser). Double stranded oligodeoxynucleotides from −1221 to −1245 of the *TLR9* promoter were purchased from Integrated DNA Technologies (Coralville, Iowa).

1. Probe with −1237C allele (C probe) 5’-AGACTTGGGGGAGTTTCCAGGCAGA-3’ 3’-TCT GAACCCCCTCAAAGGTCCGTCT-5’
2. Probe with −1237T allele (T probe) 5’-AGACTTGGGGGAGTTTTCAGGCAGA-3’ 3’-TCT GAACCCCCTCAAAAGTCCGTCT-5’

Nuclear extracts (10 μg) were incubated with 10 fmol of probe labelled with biotin using the Biotin 3’ End DNA labeling kit (#89818, ThermoScientific). Binding reactions were performed at room temperature for 20 min in 20 μL reaction buffer (total volume) containing 1X binding buffer, glycerol 2.5%, MgCl2 (5 mM), poly dI.dC (50 ng/μL), 0.05% NP-40. For competition assays, 200-fold excess of unlabeled DNA probe (cold probe) (2 pmol) was added to the reaction. Supershift assays were performed by incubating nuclear proteins (10 μg) with 0.2 or 0.4 μg/μL of NFκB antibody for 1h at room temperature before the addition of the biotin-labeled probe. Reactions were terminated by adding 5 μL of 5X loading buffer and resolved on a 6% DNA-retardation gel (EC6365BOX, ThermoFisher) at 100 V for 1h in 0.5X TBE buffer (# 28355, ThermoFisher). DNA-protein complexes were transferred to Hybond-N+ nylon membrane (# RPN303B, Amersham Pharmacia Biotech, Piscataway, NJ) on a Transblot apparatus (BioRad, Hercules, CA) in ice-cold 0.5X TBE buffer for 1 h at 100 V. After transferring, the membrane was cross-linked for 15 min on UV transilluminator equipped with a light bulb emitting 312nm. After the crosslinking, the membrane was blocked for 15 min and then incubated in blocking buffer containing stabilized Streptavidin-Horseradish Peroxidase Conjugate (66.7 Cl/20 mL blocking buffer) for an additional 15 min. The blocked membrane was then washed 4 times in 1X wash buffer for 5 min each and then incubated in Substrate Equilibration buffer for 5 min, followed by a 5 min incubation in 10 mL of Luminol enhancer solution. After the final incubation step, the membrane was exposed to X-ray film for 1-15 min.

## Supplementary Figure Legend

**Figure S1.**
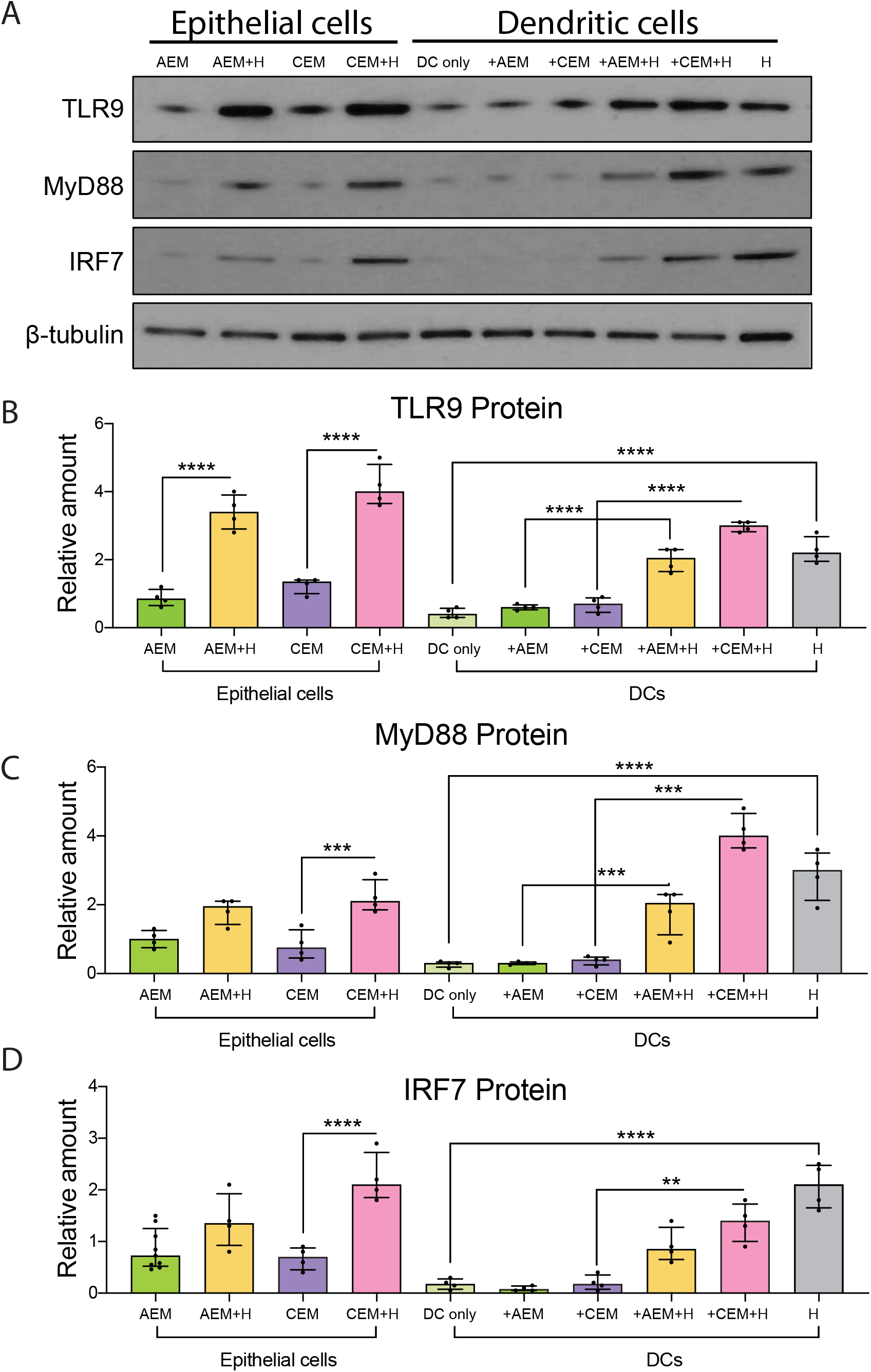
TLR9 signaling was activated by *H. felis* infection in both epithelial cells and pDCs. (A) Western blot of TLR9, MyD88 and IRF7 expression in 2D gastric epithelial monolayers derived from organoids (epithelial cells) and pDCs 48 h after initiating the co-culture with *H. felis*. Bar graphs show quantitation of western blots for (B) TLR9 (C) MyD88 (D) IRF7 using ImageJ. β-tubulin served as loading control. Horizontal lines represent the median and interquartile range for N=4 expts. \*\**P*<0.01. *** *P*<0.001. \*\*\*\**P*<0.0001.

**Table S1.** Association between TLR9 −1486T/C frequency with gastric metaplasia or gastric cancer in Vietnamese and Chinese patients.

| SNP -1486 | Vietnamese |  |  |  | Chinese |  |
| --- | --- | --- | --- | --- | --- | --- |
|  | *Hp-* G | *Hp+* G | *Hp+* A | *Hp+* Met | Healthy | GAC |
| TT | 10 | 10 | 11 | 7 | 8 | 18 |
| TC | 13 | 12 | 13 | 14 | 11 | 31 |
| CC | 5 | 7 | 6 | 6 | 3 | 14 |
| Odds Ratio<br>(95% CI) | 1 | 1.17<br>(0.57-2.41) | 1.03<br>(0.50-2.10) | 1.33<br>(0.64-2.83) | 1 | 1.30<br>(0.64-2.58) |
| C Freq (%) | 0.410 | 0.448 | 0.417 | 0.481 | 0.404 | 0.468 |
| P Value |  | NS | NS | NS |  | NS |

NS: Not significant.

## Notes

**Grant Support:** Supported by R01 DK118563 (to JLM), Arizona Comprehensive Cancer Center P30 CA023074, U54 CA143924; R01 DK083402-06A1 (YZ) and 1U19AI116491-01 grant (YZ).

### Competing Interest Statement

The authors have declared no competing interest.

